# A Model of High-Speed Endovascular Sonothrombolysis with Vortex Ultrasound-Induced Shear Stress to Treat Cerebral Venous Sinus Thrombosis

**DOI:** 10.1101/2022.11.02.514936

**Authors:** Bohua Zhang, Huaiyu Wu, Howuk Kim, Phoebe J. Welch, Ashley Cornett, Greyson Stocker, Raul G. Nogueira, Jinwook Kim, Gabe Owens, Paul Dayton, Zhen Xu, Chengzhi Shi, Xiaoning Jiang

**Affiliations:** Department of Mechanical & Aerospace Engineering, North Carolina State University, Raleigh, NC; George W. Woodruff School of Mechanical Engineering, Georgia Institute of Technology, Atlanta, GA; Department of Biomedical Engineering, University of Michigan, Ann Arbor, MI; Department of Neurology, University of Pittsburgh Medical Center, Pittsburgh, PA; Department of Biomedical Engineering, University of North Carolina, Chapel Hill, NC; Parker H. Petit Institute for Bioengineering and Bioscience, Georgia Institute of Technology, Atlanta, GA

## Abstract

This research aims to demonstrate a novel vortex ultrasound enabled endovascular thrombolysis method designed for treating cerebral venous sinus thrombosis (CVST). This is a topic of significant importance since current treatment modalities for CVST still fail in as many as 20-40% of the cases and the incidence of CVST has increased since the outbreak of the COVID-19 pandemic. Compared with conventional anticoagulant or thrombolytic drugs, sonothrombolysis has the potential to remarkably shorten the required treatment time owing to the direct clot targeting with acoustic waves. However, previously reported strategies for sonothrombolysis have not demonstrated clinically meaningful outcomes (e.g., recanalization within 30 minutes) in treating large, completely occluded veins or arteries. In this paper, we demonstrated a new vortex ultrasound technique for endovascular sonothrombolysis utilizing wave-matter interaction-induced shear stress to enhance the lytic rate substantially. Our *in vitro* experiment showed that the lytic rate was increased by at least 64.3 % compared with the nonvortex endovascular ultrasound treatment. A 3.1 g, 7.5 cm long, completely occluded *in vitro* 3D model of acute CVST was fully recanalized within 8 minutes with a record-high lytic rate of 237.5 mg/min for acute bovine clot *in vitro*. Furthermore, we confirmed that the vortex ultrasound causes no vessel wall damage over *ex vivo* bovine veins. This vortex ultrasound thrombolysis technique potentially presents a new life-saving tool for severe CVST cases that cannot be efficaciously treated using existing therapies.

## Introduction

Cerebral venous sinus thrombosis (CVST) is a pathologic blood clot formation in the cerebral venous sinuses, one of the most prevalent causes of stroke in young individuals^1,2^. The incidence of CVST has been reported as being between 2 and 13 per million per year in Europe^3–5^. More recent research reveals that the incidence of CVST is growing in the United States^6^. Over the last decade, there has been mounting evidence that early diagnosis and anticoagulant therapy minimize the morbidity and mortality associated with CVST^7^. It is common for CVST to lead to a breakdown of the blood-brain barrier and a drop in cerebral perfusion pressure, which results in cerebral edema, local ischemia, and in some cases, intracerebral hemorrhage (ICH)^1^. Although most patients respond favorably to existing treatments, some individuals fail to recover or continue to worsen despite receiving the best available treatments. The death rate for CVST patients remains at approximately 10%, with yet another 10% of individuals receiving poor long-term prognoses^1^. Moreover, in contrast to most other types of strokes, which typically occur in older people, CVST patients are much younger than the general population and are frequently pregnant women or young mothers who have recently given birth to a baby^1,8,9^. It is worth noting that a statistically significant rise in the occurrence of CVST has indeed been reported during the COVID-19 outbreak because of both the SARS-CoV-2 disease and the consequence of some vaccinations^10–18^.

Currently, the three most common therapeutic options for CVST are systemic use of anticoagulants or thrombolytic medicines, decompressive craniectomy, and catheter-based endovascular therapy (EVT)^1,19,20^. According to a recent randomized study, long-term anticoagulation failed to recanalize 33% and 40% of the patients receiving intravenous heparin accompanied by warfarin and dabigatran, respectively, despite the use of long-term anticoagulation for at least five months^21^. Decompressive craniectomy is a very invasive neurosurgical intervention consisting of partial skull removal, which although potentially lifesaving, it is often considered the last resource since it decompresses the edematous brain without addressing the underlying problem (e.g., occlusive venous clot)^19^. Thrombolytic medications such as tissue plasminogen activator (t-PA) destroy blood clots by dissolving the cross-linked fibrin proteins that form the structure of the clots^22^. Nevertheless, systemic fibrinolytic medications are frequently inefficient, require extended treatment periods (up to 16 hours), and may result in severe hemorrhage in as many as 55% of patients, with around 10% of those instances resulting in possibly fatal ICH^1,23–26^. There have been a variety of catheter-based devices that may be used to treat clots in the target area, such as mechanical thrombectomy and catheter-directed administration of thrombolytic medicines. In severe CVST instances, mechanical thrombectomy is becoming more popular^27^.

However, existing endovascular techniques are insufficiently successful because they are not specially intended to treat the venous sinus, which has an average diameter of more than three times greater than that of the intracranial arteries. Furthermore, those techniques provide a high probability of severe consequences, including vascular endothelial damage, which might result in deadly ICH^24,28^. When thrombolytic medications are delivered locally by catheter, the risks associated with systemic administration are reduced, and the drugs are more successful at delivering into the clot region. Despite this, the effectiveness of this approach is uncertain, and it is still associated with the concerns of ICH^22,23,28,29^. According to a newly published randomized clinical study, the inadequate efficiency of existing device technology is vividly highlighted by a limited group of participants (n=33) compared to systemic anticoagulation (n=34). Although EVT has a favorable safety record, the trial could not establish a therapeutic advantage^30^. The fact that preliminary randomized studies using the early thrombectomy technique failed to establish its effectiveness in arterial strokes, whereas later studies with second-generation arterial thrombectomy were resoundingly beneficial is noteworthy^31,32^. It is thus acceptable to speculate that, with new technical improvements, venous thrombectomy may one day be demonstrated to be advantageous in properly selected patients.

In order to overcome these constraints and enhance effective thrombus breakdown without raising the danger of cerebral or systemic hemorrhage consequences, ultrasound thrombolysis, also known as sonothrombolysis, has been explored^33^. The primary mechanisms involved in sonothrombolysis are the radiation force, acoustic streaming, and cavitation^34^. It has been discovered that fibrinolytic medications and contrast agent-mediated ultrasound may speed up thrombolysis by increasing the transport of drugs into the clot^34–37^. To improve the clot lysis efficiency, large aperture transducers were employed in conjunction with multi-frequency excitations to maximize the acoustic cavitation effect^34,38–40^. Using a catheter-based ultrasound transducer (EKOS), a multi-center retrospective study found that ultrasound combined with t-PA shortened infusion time^41–43^ and resulted in a higher rate of completely dissolving clots for deep venous thrombosis therapy^44,45^. Sonothrombolysis efficacy has been improved using microbubbles (MBs) and nanodroplets (NDs) combined with forward-viewing transducers^46,47^. Sonothrombolysis is therefore a promising therapy for the treatment of deep vein thrombosis. However, there is currently no endovascular ultrasound catheter that can be used for CVST treatment because of the extended treatment period (> 15 hours) and the high t-PA dosage used (10-20 mg)^48^ as well as the challenges related to the navigation into the more tortuous intracranial vessels. In addition, conventional sonothrombolysis therapies, mainly based on the mechanism of cavitation effects, often require a high peak negative pressure, which may be unsafe and challenging to accomplish in endovascular sonothrombolysis^49^. In this context, there is an unmet medical need for a thrombolysis strategy that can provide local, effective, and rapid lysis of various types of clots, such as large acute clots and completely occluded clots, while minimizing damage to the vessel and surrounding tissue, as well as many other medical symptoms related to high doses of drugs for CVST care.

Vortex ultrasound (also known as acoustic orbital angular momentum) is a form of the acoustic wave that propagates across space with a helical wavefront that rotates as it moves through the space^50^. The vortex ultrasound-induced shear force has the potential to break down clots safely and improve the efficacy of thrombolysis. The objective of this work is to demonstrate the endovascular vortex ultrasound (EVUS) thrombolysis (Fig. 1a) for safe and effective CVST treatment in order to achieve a significant breakthrough in addressing the unmet medical sonothrombolysis challenges mentioned above^46,47^. A 2 by 2 array of four small aperture, low-frequency (1.8 MHz) piezoelectric transducers (Fig. 1b), each with a forward-viewing surface that is shifted by a quarter wavelength along the wave propagation direction, can be patterned in this manner to produce a physical helical wavefront (Fig. 1c). The transducer array was assembled into a 9-French catheter (with the diameter of about 3.0 mm) combined with a lumen for cavitation agents and drug delivery (Fig. 1b). This study demonstrates for the first time that the vortex ultrasound-mediated by the microbubbles induces localized shear stress to the blood clot (Fig. 1d) that is expected to increase the sonothrombolysis rate significantly. This new EVUS system enables effective and rapid treatment of large acute and fully occluded clots, thereby significantly reducing damage to the vessel and surrounding tissue and shrinking the size of clot debris, reducing the risk of recurrent and distal embolisms.

**Fig. 1.**
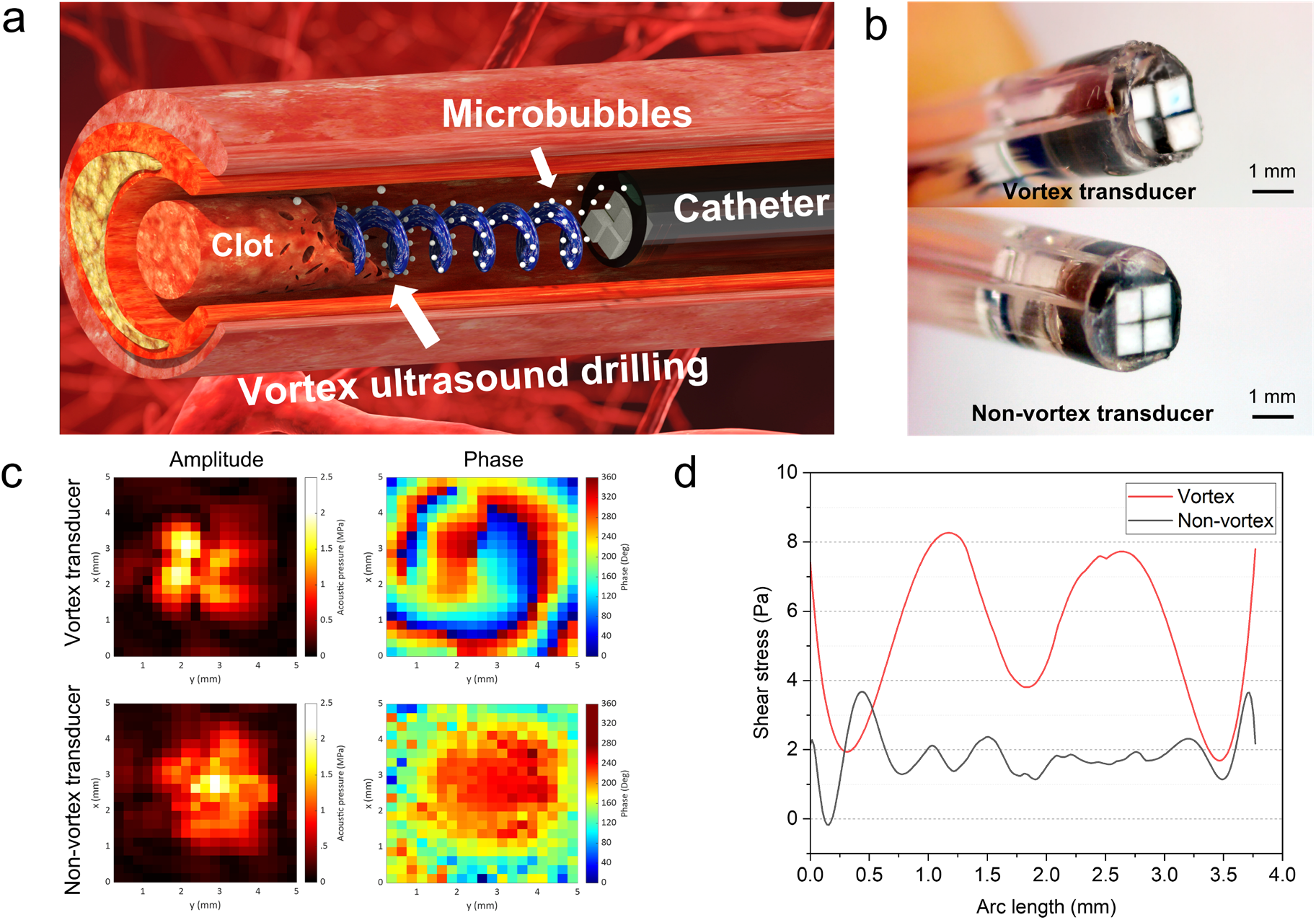
High-speed endovascular sonothrombolysis with vortex ultrasound-induced shear stress. **a**, The schematic view of the sonothrombolysis process with a vortex ultrasound transducer. The vortex ultrasound transducer is installed in a 9-Fr catheter and inserted into the blood vessel with a blood clot. The microbubble cavitation agents are injected through a drug delivery lumen of the catheter. The blood clot breaks up under the shear stress and cavitation effects of vortex ultrasound. **b**, The prototype of the developed vortex transducer and nonvortex transducer installed in a 9-Fr catheter with drug delivery lumen. **c**, The measured acoustic pressure map for vortex and nonvortex transducer. The phase map of the vortex transducer shows the swirling shape, and the nonvortex transducer only shows one circular shape. **d**, The COMSOL calculated shear stress distribution in the blood clot along the azimuthal direction under the exposure of vortex and nonvortex ultrasound stimulation. The vortex ultrasound induces about four-fold greater peak shear stress inside the blood clot than the nonvortex ultrasound.

## Results

### Vortex ultrasound transducer

In this study, we leveraged the multilayer forward-viewing ultrasound transducer technology to develop a vortex ultrasound transducer array (Fig. 1b) to generate the helical wavefront (Fig. 1c). The four transducers were attached to an epoxy base containing air bubbles with a quarter wavelength (0.21 mm) step between neighboring transducers to form a 2 by 2 helical-patterned transducer array (Supplementary Fig. 1a) for vortex ultrasound generation. Each transducer had an aperture of 0.8 × 0.8 mm^2^ and a longitudinal-excitation-mode resonance frequency of 1.8 MHz. Air bubbles were introduced into the epoxy substrate to improve the acoustic contrast on the rear sides of the transducers and the forward ultrasonic emission. The transducer array prototype had an overall aperture of about 1.65 × 1.65 mm^2^. For microbubble and lytic agent distribution, a 9-French 2-lumen catheter was used in conjunction with a duct channel explicitly designed for this purpose. Using a calibrated hydrophone, we measured the emitted acoustic pressure field of the prototyped transducer array. A pressure field was generated using a 60 V_pp_ voltage input and was measured both in amplitude and phase at a distance of approximately 1.5 wavelengths (1.2 mm) away from the transducer array (Fig. 1c). Noticeably, the pattern in the pressure amplitude was toroidal-like, as predicted for an acoustic vortex beam, and a spiral pattern was observed in the pressure phase. The vortex ultrasonic beam (insonation zone) diameter at -6 dB is about 2.3 mm, and it is predicted to be broader downstream due to diffraction as it propagates. The peak mechanical index (MI), a metric of measuring ultrasound bioeffects defined by the ultrasound beam’s peak negative pressure (PNP) divided by the square root of the operating frequency, achieved in the near field (0.5 wavelengths) of the transducer array is approximately 1.5, below the FDA’s MI limit of 1.9^51^. It is expected to be approximately 0.9 at about 1.5 wavelengths, which is sufficient to cause microbubble-mediated cavitation for enhanced sonothrombolysis while remaining safe for intravascular operation^46,47^. Compared to vortex ultrasound’s spiral propagation pattern, an array of transducers with a flat front viewing surface generates a standard Gaussian beam profile (Fig. 1c), with the peak pressure amplitude located in the middle of the beam.

One of the most prominent advantages of vortex ultrasound is the strong in-plane pressure gradient that creates a rotational shearing stream in fluids^52^ and considerable shear stress in the interacting solids when applied^53,54^. For rapid and safe CVST therapy, the induced shear stress in blood clots loosens and breaks the fibrins, increasing the sonothrombolysis rate and decreasing the necessary treatment time and lytic agent dosage. Numerical calculations performed using COMSOL Multiphysics revealed that the vortex ultrasound transducer induced larger shear stress along the azimuthal direction than conventional plane waves produced by nonvortex ultrasound transducer (Fig. 1d). Based on the simulation results, the Reynolds shear stress of the shear flow is around 80 dyne/cm^2^, which is much lower than the lowest recorded hemolysis threshold 2,500 dyne/cm^2^ in magnitude^55^, which is confirmed by our hemolysis tests (Supplementary Fig. 8). Thus, the shear stress induced by the vortex ultrasound has no potential to cause damage to the blood cells.

### In vitro vortex sonothrombolysis

Because CVST is often associated with clots that are just a few hours to a few days old^56^, termed acute to subacute clots, our *in vitro* studies were performed to determine the rate of sonothrombolysis of acute clots using the prototyped vortex ultrasound transducer. A comparison of *in vitro* thrombolysis therapy outcomes between nonvortex and vortex ultrasound transducer treatment was shown in Fig. 2. The results illustrate the clot reduction during the treatment in 5 min intervals (Fig. 2a), and the vortex transducer shows a significantly higher clot lysis speed than the nonvortex transducer for 30 minutes of sonothrombolysis treatment. Following the treatment, the residual blood was washed out with saline (Fig. 2b) to compare the results of nonvortex and vortex ultrasonic treatments performed at the same push-through force level (0.056± 0.021 N, feed-in speed: 1.66 mm/min). As indicated by the horizontal white dashed lines, the vortex transducer recanalized the whole length of the 50 mm acute clot, but the nonvortex transducer achieved less than 50% of clot lysis within the 30-minute treatment period and did not recanalize the blood vessel phantom (Fig. 2a).

**Fig. 2.**
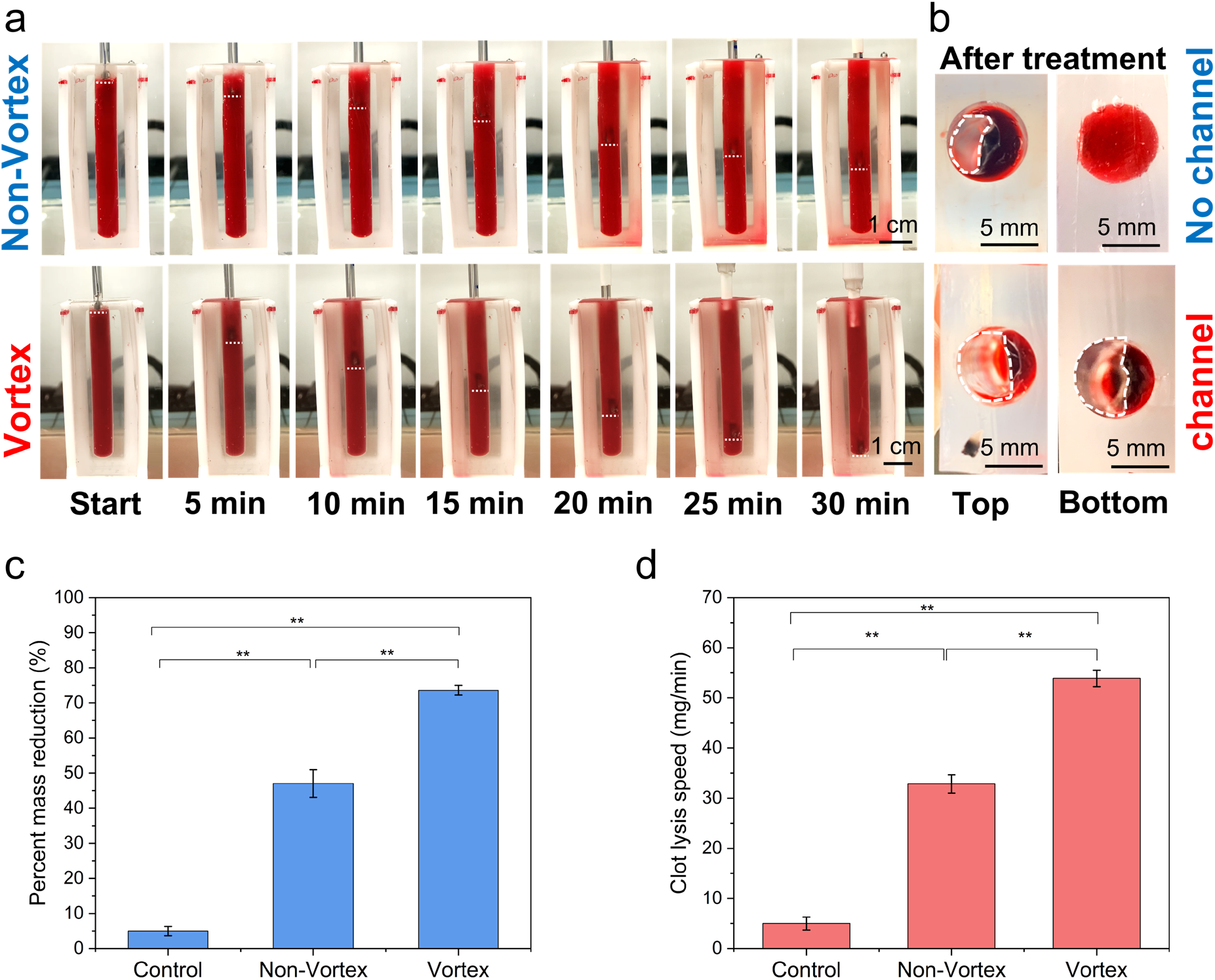
The *in vitro* thrombolysis treatment results comparison for nonvortex and vortex ultrasound transducer treatments. **a**, 30 min treatment compared with the same level of push-through force. The horizontal white dashed lines mark the current location of the ultrasound transducer. During the 30 min treatment process, the vortex transducer exhibited a significantly higher clot lysis speed than the nonvortex transducer. **b**, After 30 min treatment, the nonvortex transducer only treated about half of the blood clot (top channel opening size: 10.2 ± 0.7 mm^2^), whereas the vortex ultrasound transducer formed a channel (flow channel opening size: 23.5 ± 0.8 mm^2^) through the entire blood clot. **c**, The percentage of the blood clot mass reduction comparison between the nonvortex and vortex transducer treatments. **d**, The comparison of blood clot lysis speed between the nonvortex and vortex transducer treatments. All the control group was only injected with saline without ultrasound treatment. (** p<0.01, n=3)

Moreover, the top and bottom cross-sections of blood clots (Fig. 2b) from the vortex treatment showed that the vortex transducer created an opening in the clot with a width of 3.6 ± 0.3 mm. In order to determine the clot lysis rate for each case, we measured the clot mass both before and after the treatment. For the percent mass reduction of the clot, the nonvortex transducer had a lysis rate of 1.57 %/min, and the lysis rate of vortex transducer-based thrombolysis was measured to be about 2.45 %/min, showing a significant increase of 1.56-fold rate over the nonvortex transducer (Fig. 2c). Comparing clot lysis speed, the nonvortex transducer-based thrombolysis yielded an absolute lysis rate of 32.8 mg/min. In contrast, the vortex transducer had an absolute lysis rate of 53.9 mg/min, suggesting a significant increase of 64.3% with the vortex sonothrombolysis (Fig. 2d).

### In vitro parameter optimization

Next, we performed the *in vitro* parameter study to optimize the ultrasound input parameters for efficient sonothrombolysis with a vortex ultrasound transducer (Fig. 3). First, the different input voltages were used as the input factor for the vortex ultrasound transducer treatment. The result showed that the percent mass reduction of the clot significantly increased from 33.7 % to 85.6 % with the increasing input voltage from 20 V_pp_ to 100 V_pp_ (Fig. 3a). Additionally, the clot lysis speed significantly increased from 23.5 mg/min to 59.7 mg/min as the input voltage increased from 20 V_pp_ to 100 V_pp_ (Fig. 3d). Our earlier study demonstrated that greater input voltages could result in higher PNP and MI^46^. However, in order to keep the ultrasound transducer operating within a safe voltage range, 100 V_pp_ was applied as the maximum input voltage due to the ultrasound transducer material limitation (45% of the AC depoling voltage for PZT-5A ceramics). For the safety of the treatment^57^, 80 V_pp_ was selected as the optimal input voltage for this study.

**Fig. 3.**
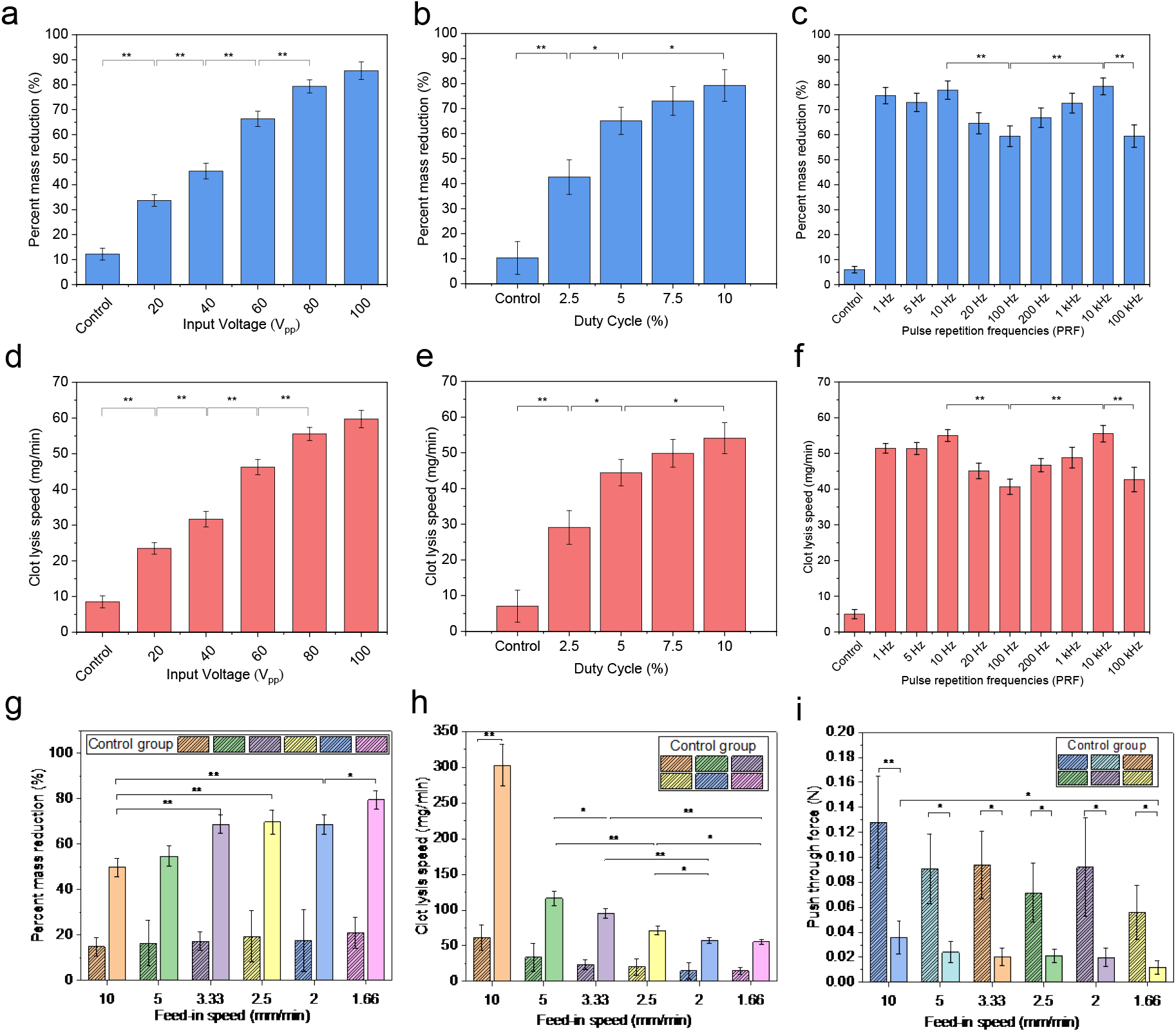
The parameter study of *in vitro* sonothrombolysis with vortex ultrasound transducer treatment. **a**, The percentage of blood clot mass reduction increases as input voltage increases from 20 V_pp_ to 100 V_pp_. **b**, The percentage of blood clot mass reduction increases as the duty cycle increases from 2.5% to 10%. **c**, The percentage of blood clot mass reduction varies with different pulse repetition frequencies (PRF). The percent mass reduction rate is highest when PRF is 10 Hz and 10,000 Hz. **d**, The lysis speed of blood clots increases with increasing input voltage. **e**, The blood clot lysis speed increases as duty cycle increases from 2.5% to 10%. **f**, The lysis speed of blood clots changes with different pulse repetition frequencies (PRF). **g**, The percentage of blood clot mass reduction changes with different catheter feed-in speeds and is significantly higher than controls across all feed-in speeds tested. **h**, The lysis speed of blood clots changes with different catheter feed-in speeds. **i**, The measured blood clot push-through force with different catheter feed-in speeds. The solid bar data present the vortex ultrasound treatment group. In contrast, the dashed bars denote the control group, which was only injected with saline without vortex ultrasound treatment. (* p<0.05, ** p<0.01, n=3)

Different duty cycles were tested (under same PRF but different burst cycles) as the next input factor for optimizing vortex ultrasound transducer treatment. The results illustrated that the percent mass reduction of the clot significantly increased from 42.6% to 79.2% when the duty cycles increased from 2.5% to 10% (Fig. 3b). The clot lysis speed also increased considerably from 29.1 mg/min to 54.1 mg/min as the duty cycles increased from 2.5 % to 10 % (Fig. 3e). Since higher duty cycles (≥10%) may induce additional safety risks like heating effects^58^, a duty cycle at 7.5 % was selected as the optimal duty cycle for this study.

Third, we tested different pulse repetition frequencies (PRF) (under same duty cycles but different burst cycles) to optimize this input factor. The results demonstrate that the percent mass reduction of the clot first significantly decreased from 77.8 % to 59.4 % with the increase of PRF from 10 Hz to 100 Hz and later significantly increased from 66.8 % to 79.3 % with the increase of PRF from 200 Hz to 10 kHz, but then drops dramatically to 59.5 % when PRF further increased to 100 kHz (Fig. 3c). Similar trends can be found in the clot lysis speed when the PRF is varied from 10 Hz to 100 kHz (Fig. 3f). The 10 Hz and 10 kHz cases outperformed other cases, which was possibly correlated with the timing of microbubbles’ intact traveling, oscillations, and ruptures. We selected 10 kHz for further tests since a previous study showed that the combination of “short-burst cycles and higher PRF” suppressed ultrasound-induced heating compared to the combination of “long-burst cycles and lower PRF” while maintaining the same duty cycle^58^.

Next, the different feed-in speeds were tested as the input factor. Feed-in speed is the rate at which the catheter was pushed through the occluded vessel. This study revealed that the percent mass reduction of the clot significantly increased from 49.8% to 79.6% when feed-in speed decreased from 10 mm/min to 1.66 mm/min (Fig. 3g). However, the clot lysis speed significantly decreased from 302.9 mg/min to 55.4 mg/min as the feed-in speed decreased from 10 mm/min to 1.66 mm/min (Fig. 3h). Although higher feed-in speed would significantly increase the clot lysis speed, it may also cause safety issues such as the large clot debris induced by fast mechanical push-through force. Therefore, the push-through force was measured under different feed-in speeds to determine the optimized parameter. These experiments demonstrated that the push-through forces with vortex ultrasound were about 4 times lower than the control group (without ultrasound) with different feed-in speeds (Fig. 3i), which proved the effectiveness of vortex ultrasound treatment. By overall consideration of the percent mass reduction (Fig. 3g) and clot lysis speed (Fig. 3h) while minimizing the induced push-through force (Fig. 3i), the feed-in speed of 3.33 mm/min was selected as the optimal parameter for this study.

### Cerebral venous sinus 3D model

An *in vitro* cerebral venous sinus 3D phantom flow model with an average sinus diameter of 10 mm was used to test the performance of the vortex transducer in treating CVST. The results showed that the completely occluded blood vessel was recanalized in only 8 minutes of treatment with a vortex ultrasound transducer (Fig. 4 and Supplementary Video 1). The measured clot mass before (3.1 ± 0.3 g) and after (1.2 ± 0.4 g) the treatment indicated that the vortex ultrasound transducer could achieve a high clot mass reduction rate (7.66 %/min) and clot lysis speed (237.5 mg/min) in 8 min treatment, which is the significantly higher thrombolysis rate than the recent tPA-free endovascular sonothrombolysis (1.3-2.5 %/min, 2-4.6 mg/min)^46,47,59–61^. Moreover, the clot debris analysis revealed that most of the clot debris particle sizes were less than 100 µm (Supplementary Fig. 5), which indicated a low risk of dangerous embolus formation. The histology results of the bovine blood vessel wall cross-sections after operating the catheters of the vortex, nonvortex, and control groups (Supplementary Fig. 6b, 6c, 6d, respectively) and pixel comparison results (Supplementary Fig. 6e) confirmed the safety of the surrounding vessel and tissues of the vortex ultrasound treatment.

**Fig. 4.**
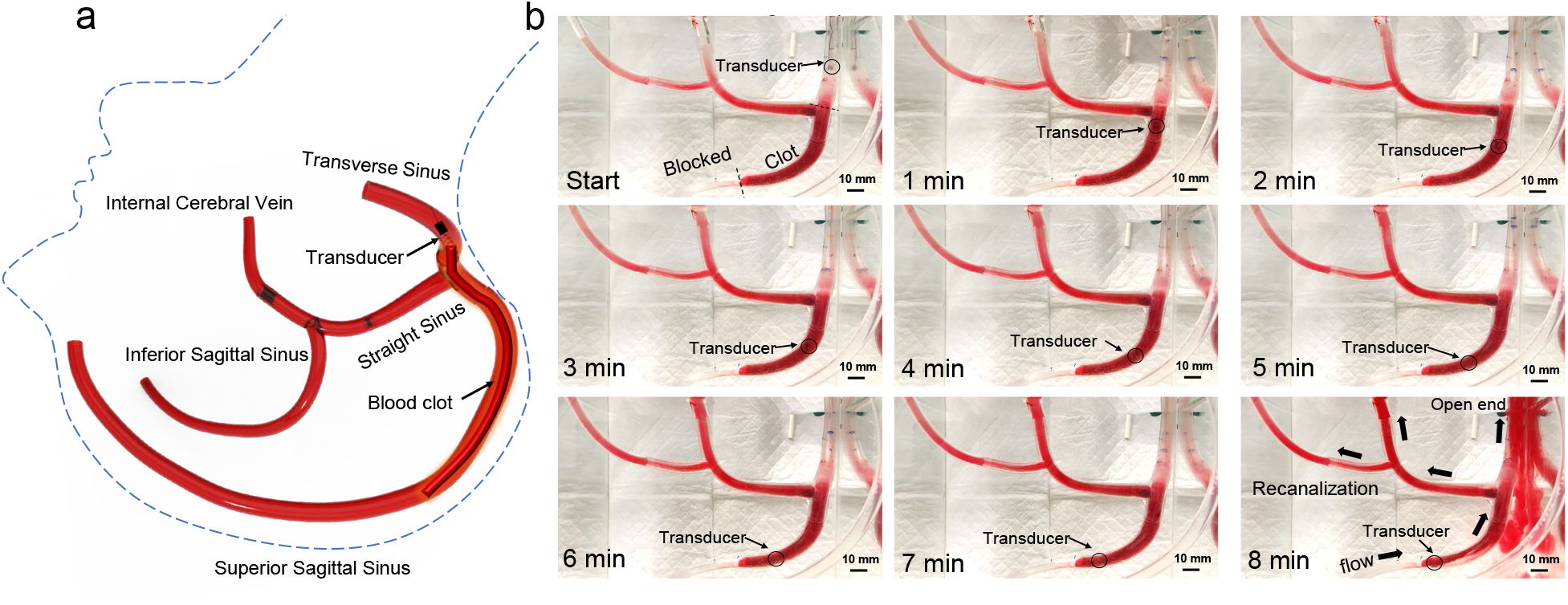
The 3D phantom study of *in vitro* sonothrombolysis with vortex ultrasound transducer treatment. **a**, Diagram of the cerebral venous sinus 3D phantom model with blood vessels labeled as well as the position of the blood clot, which runs along the superior sagittal sinus into the transverse sinus. The transducer starts in the transverse sinus. An outline of a human head and neck are included in the diagram, as indicated by the blue dashed line, to orient the position of these vessels *in vivo*. **b**, A time lapse of the blood clot treatment (clot length: 75 ± 3 mm, clot age: 1 hour, treatment time: 8 minutes, vortex ultrasound transducer input factors: 80 V_pp_ operating voltage, 7.5 % duty cycle, frequency= 1.8 MHz, PRF=10 kHz) (scale bar = 10 mm). The blood flow direction is marked by the thick black arrows in the final panel. The flow is driven by a pressure applied before and during the treatment to mimic the *in vivo* environment. The location of the vortex transducer catheter tip is marked by the black circle in each panel.

## Discussion

Here, we demonstrate a novel EVUS system with a small aperture array that generates vortex ultrasound with a helical pattern to induce localized shear stress in the blood clot, which dramatically accelerates sonothrombolysis and lowers the necessary drug dose and MI for clot dissolution in an *in vitro* 3D model of acute CVST. This device was the first to incorporate the novel contrast agent-mediated vortex ultrasound for clot-dissolving technology into a 9-French catheter device to demonstrate the remarkably increased lytic efficiency and safety over existing thrombolysis approaches.

There is no endovascular thrombolysis technology currently available to treat severe CVST effectively. Therefore, this study developed a unique endovascular forward-viewing ultrasound transducer array with a helical pattern to generate vortex ultrasound in the cerebral venous sinus. For illustration, a 2 by 2 array of four multilayer stacked transducers (Supplementary Fig. 1a) with resonance frequency at 1.8 MHz, with the forward viewing surfaces of neighboring transducers shifted by a quarter wavelength, has been used to induce a physical ultrasound phase delay, which is required to generate the helical wavefront of vortex ultrasound.

The vortex ultrasound produces an in-plane pressure gradient, which causes localized shear stress in the blood clot. The localized shear stress and cavitation-induced (Supplementary Fig. 7) microstreaming significantly accelerate fibrinolysis in the clot, increasing the sonothrombolysis rate while simultaneously decreasing the medication dosage, peak-negative pressure (PNP), and treatment duration. The shear stress induced by the vortex ultrasound loosens the clot structure, which improves the delivery of MBs and the lytic agent into the CVST. It should be pointed out that the induced shear stress would not present a hemolysis concern since the Reynolds shear stress of the shear flow is approximately 80 dynes/cm^2^ from the simulation, which is significantly less than the lowest known hemolysis threshold (2,500 dynes/cm^2^)^55^. Besides, the *in vitro* hemolysis test results showed that the plasma-free hemoglobin (PfHgb) level was about 5.7 to 23.4 mg/dL for 5 to 30 minutes treatment with vortex ultrasound and microbubbles (Supplementary Fig. 8), which was significantly lower than the clinical signs of hemolysis (>40 mg/dL)^62^ and treatment results from other technologies such as OmniWave (228 mg/dL), histotripsy (348 ± 100 mg/dL)^63^ and AngioJet (1367 mg/dL)^64^. For the first time, a low-frequency (1.8 MHz), safe (MI 0.5-1.5) vortex ultrasound beam was employed to achieve quick and safe catheter-directed CVST sonothrombolysis using a vortex ultrasound beam. Therefore, this study developed a unique, highly efficient, and safe sonothrombolysis approach as preclinical research, which will fulfill the therapeutic demands of patients with totally blocked, massive CVSTs in the future.

In summary, we described a novel vortex transducer technology specifically designed for the treatment of CVST. The developed vortex transducers showed an absolute lysis rate of 53.9 mg/min without parameter optimization, which is 64.3 % higher than the lysis rate of nonvortex transducers (32.8 mg/min) at the same push-through force level. For the optimized case, the improvement is expected to be even more significant. We demonstrated that the vortex ultrasound transducer could potentially fully recanalize the completely occluded acute CVST *in vitro* within 8 minutes of treatment and achieve a record-high clot lysis speed (237.5 mg/min). In severe cases of CVST and in patients with massive, fully blocked venous clots and who cannot be effectively treated with medications that are currently available, the vortex ultrasound thrombolysis technology may become a life-saving treatment in the future. Additional studies using a novel CVST animal model are planned.

## Methods

### Transducer design

The azimuthal polar coordinate of each transducer element is used to construct the vortex ultrasound transducer array to establish the suitable acoustic phase delay of each transducer element. The acoustic phase delay (*φ*) of an element with an in-plane azimuthal polar coordinate *θ* is provided by *φ* = *lθ* to create vortex ultrasound with topological charge *l* (a quantity that represents the angular momentum carried by the vortex wave)^65,66^. Topological charges of greater magnitude suggest the presence of a vortex wave with higher angular momentum and a larger aperture^65^. The 2 by 2 transducer array design (Supplementary Fig. 1) can produce vortex ultrasound with *l* = ±1, allowing us to achieve the minimum aperture size possible with this configuration. In this scenario, the acoustic phase delay between adjoining transducers is *π*/2, which corresponds to a quarter wavelength shift in their forward viewing surfaces when compared to one another. By employing the epoxy base, it is possible to align the four components with a quarter wavelength (0.21 mm for 1.8 MHz) shift between the forward viewing surfaces of adjoining transducers to achieve the desired result.

### Transducer fabrication

The fabrication procedure for the proposed transducer arrays is shown in Supplementary Fig. 2. In this study, two piezoelectric plates (for example, PZT-5A, with an area of 6 × 6 mm^2^ and thickness of 200 μm) were bonded together using steel-reinforced epoxy (8265S, J-B Weld Company, Sulphur Springs, TX, USA) with a thickness of around 20 μm. A quarter-wavelength matching layer composed of an alumina powder/epoxy bond combination with an acoustic impedance of 5-6 MRayls was added to the front side of the device. An air bubble/epoxy composite backing was applied on the backside of the piezoelectric plates with a thickness of about 6 wavelengths (1.5 mm). By lapping with the backing layer, the aperture height of four piezoelectric multilayers that were integrated with the matching and the backing varied by a quarter wavelength. The bonded stacks were diced for an element aperture of approximately 0.8 × 0.8 mm^2^ (DISCO 322, DISCO Hi-Tec America, Inc., San Jose, CA), yielding four multilayered stacks with varying aperture heights. The multilayered stacks were bonded using an electrically nonconductive alumina/epoxy composite. After utilizing epoxy to isolate unnecessary electrodes, the transducer electrodes were connected with a coaxial cable (5381-006, AWG 38, Hitachi Cable America Inc., Manchester, NH). The piezoelectric transducers were combined into a two-lumen flexible catheter with a 9-French diameter; one lumen guided the transducer while the other served as a flow channel for administering drugs and contrast agents. The catheter was composed of polyethylene, which allowed the 9-French catheter to be flexible enough to be directed into the cerebral venous sinus during the procedure.

### Transducer characterization

Following fabrication, the impedance spectrum was examined by measuring the resonance frequency of the device. Using a calibrated needle hydrophone (HNA-0400, ONDA Corp., Sunnyvale, CA), the transducer array was installed on a computer-controlled 3-axis translational stage (Anet A8, Anet Technology Co., Ltd., Shenzhen, China) to characterize the acoustic waveform and pressure output (Supplementary Fig. 3). Using a function generator (33250A, Agilent Tech. Inc., Santa Clara, CA), a sinusoidal pulse with ten cycles every ten microseconds was sent to an RF power amplifier (75A250A, AR, Souderton, PA). The signal was amplified before being sent into the developed transducer.

### Acute clot preparation

Bovine blood was used to prepare the acute clot in a manner similar to our previous study^46,47^. Initially, anticoagulated bovine blood (Lampire Biological Laboratories, Pipersville, PA, USA) containing acid citrate dextrose (ACD) was combined with 2.75% Calcium Chloride (Fisher Scientific, Fair Lawn, NJ, USA) in a 10:1 ratio (100 mL blood/10 mL CaCl_2_) to form the blood mixture. Next, the blood mixture solution was added to the PDMS channel (diameter: 7 mm) to form the acute blood clots (length: 50 ± 3 mm, diameter: 7 ± 0.5 mm). For the 3D phantom *in vitro* study, the blood mixture solution was injected into the 3D phantom channel to form the acute clot. Finally, the acute blood clots were incubated in the water bath (PolyPro Bath, Model RS-PB-100, USA) at 37 °C for one hour before use.

### Microbubble preparation

The microbubbles were prepared in-house as described in previous studies^46,67,68^. In brief, lipid mixtures were created by combining DSPC and DSPE-PEG2000 at a 9:1 molar ratio (Avanti Polar Lipids, Alabaster, AL, USA) in a solution that also included propylene glycol at a concentration of 15% (v/v), glycerol at a concentration of 5% (v/v), and phosphate-buffered saline at a concentration of 80%. After that, aliquots of lipid solution measuring 1.5 ml each were transferred to glass vials measuring 3 ml, and the air headspace in each vial was replaced with decafluorobutane gas (Fluoromed, Round Rock, TX, USA). Microbubbles containing decafluorobutane gas cores and phospholipid shells may spontaneously develop when agitation using a Vialmix device (Lantheus Medical Imaging, N. Billerica, MA, USA) is performed. Single particle optical methods (Accusizer 780, Particle Sizing Systems, Santa Barbara, CA, USA) were used to assess the concentration of microbubbles as well as their diameter. The average size of the microbubbles was 1.1 µm, and their concentration was 10^10^/mL in each vial. The microbubbles solution was then diluted into 10^9^/mL for each *in vitro* test.

### *In vitro* test preparation

For each in vitro treatment experiment, the acute blood clot was placed inside the PDMS channel. As shown in Supplementary Fig. 4, the PDMS channel was fixed on the balance (SPX123, OHAUS, Parsippany, NJ, USA) that was connected to a computer for the push-through force measurement in real-time. A 3D motion stage was used to control the catheter feed-in speed to maintain the distance between the clot and vortex ultrasound transducer, about 1.0 mm within the focal zone. The vortex ultrasound transducer was powered by a radio frequency power amplifier (Amplify ratio: 53 dB, Model: 75A250A, AR, Inc., Souderton, PA, USA), while the input sine wave signal was generated by a function generator (Model: 33250A, Agilent Technologies, Inc., Loveland, CO, USA). The vortex and nonvortex transducers were operated with an 80 V_pp_ (peak-to-peak) input voltage, a duty cycle of 7.5%, and a pulse repetition frequency of 10 kHz. The input voltage, duty cycle, pulse repetition frequency, and feed-in speed for the optimal parameters study may vary according to the different experimental conditions. The microbubbles solution with a concentration of 10^9^/mL was delivered through the catheter using a microfluid pump (DUAL-NE-1010-US, New Era Pump Systems Inc., Farmingdale, NY, USA) with an infusion rate of 0.1 mL/min. A clot sample with a length of about 50 ± 3 mm and a weight of about 1.6 ± 0.4 g was employed for each test. The length of the remaining clots was measured every 5 minutes for a total of 30 minutes. Following the treatment, the lysis rate and speed were computed, and the channel width was evaluated in order to compare the efficacy of the vortex and nonvortex transducers in terms of channel width. For the in vitro 3D phantom experiment (Fig. 4), an acute blood clot (diameter: 10 ± 2 mm, length: 75 ± 3 mm) was incubated in the phantom channel. The 3D-printed phantom channel was filled with saline and maintained at a temperature of 37.5 ± 0.5 °C. The saline was pumped from the reservoir to the 3D phantom venous flow model using a peristaltic pump, and a valve was utilized to modify the inlet flow speed and liquid pressure. The fluid pressure was measured with a digital pressure gauge and maintained at 50.3 mmH_2_O by regulating the pumping speed and valve before the flow channel. The flowing outlet of the phantom channel was connected to a small water reservoir to collect the saline-containing clot fragments following thrombolysis treatment.

### Vessel wall damage study

To evaluate the vessel damage caused by sonothrombolysis treatment, we performed the *ex-vivo* safety tests with vortex and nonvortex transducer using the Canine jugular veins (NC State College of Veterinary Medicine, Raleigh, US). Before usage, bovine leg veins were kept in cold PBS, and then the vessels were cleaned and trimmed to approximately 1 cm in length (Supplementary Fig. 5a). After the split and mounted in a water tank with degas water, samples were treated in 3 groups with nonvortex ultrasound + MBs and vortex ultrasound + MBs, respectively. Each test takes 30 mins with the standard ultrasound parameters for the sonothrombolysis. Another untreated sample from the same jugular vein was picked as the control group. The treated vessels were then clipped to isolate the region of excitation and fixed in formalin for histological examination.

Hematoxylin and eosin (H&E) staining was performed for the three groups of samples. Three mirrored cross-sections were obtained for the sample at 100 μm (Supplementary Fig. 5). After that, the slides were scanned using the EVOS FL Auto system (EVOS FL Auto Imaging System, Life Technologies Corporation, Carlsbad, USA) with 20X magnification. For each group, 6 regions were randomly selected along with the cross-section view, and different pixels were picked with a binary threshold. The pixel ratios representing the red color and white color were averaged and compared for the three groups.

### Clot debris study

To evaluate the clot debris induced by sonothrombolysis treatment, we collected the blood solution contained with clot debris after each treatment. The blood solution was filtered with nylon plastic mesh sizes 100 and 50 μm (Supplementary Fig. 6). After being dried for at least 24 hours, the meshes were examined under the microscope, and the clot debris particle size was estimated by ImageJ software (Rasband WS, ImageJ, U. S. National Institutes of Health, Bethesda, MD, USA). A one-way analysis of variance (ANOVA) was conducted for statistical significance in clot particle diameter and variances, with a significance level of 0.05 used to determine statistical significance.

## Author contributions

BZ, HK, HW, ZX, CS, and XJ designed the research.

BZ, HK, HW, AC, and GS performed the experiments.

BZ, HK performed the simulations.

BZ, XZ, GO, CS, and XJ analyzed the data.

BZ. wrote the original draft.

BZ, HW, PJW, JK, ZX, CS, and XJ wrote and edited revised drafts.

CS, XJ supervised the study.

CS, XJ, PD, ZX acquired the funding.

All authors provided active and valuable feedback on the manuscript.

