## Supplementary Materials for "A Model of High-Speed Endovascular Sonothrombolysis with Vortex Ultrasound-Induced Shear Stress to Treat Cerebral Venous Sinus Thrombosis"

Bohua Zhang[1,*], Huaiyu Wu[1,*], Howuk Kim[1,*], Phoebe J. Welch[2], Ashley Cornett[3], Greyson Stocker[3] Raul G. Nogueira[4], Jinwook Kim[5], Gabe Owens[3], Paul Dayton[5], Zhen Xu[3], Chengzhi Shi[2,6,†], Xiaoning Jiang[1,†]

[1]Department of Mechanical & Aerospace Engineering, North Carolina State University, Raleigh, NC

[2]George W. Woodruff School of Mechanical Engineering, Georgia Institute of Technology, Atlanta, GA

[3]Department of Biomedical Engineering, University of Michigan, Ann Arbor, MI

[4]Department of Neurology, University of Pittsburgh Medical Center, Pittsburgh, PA

[5]Department of Biomedical Engineering, University of North Carolina, Chapel Hill, NC

[6]Parker H. Petit Institute for Bioengineering and Bioscience, Georgia Institute of Technology, Atlanta, GA

[*]These authors contributed equally

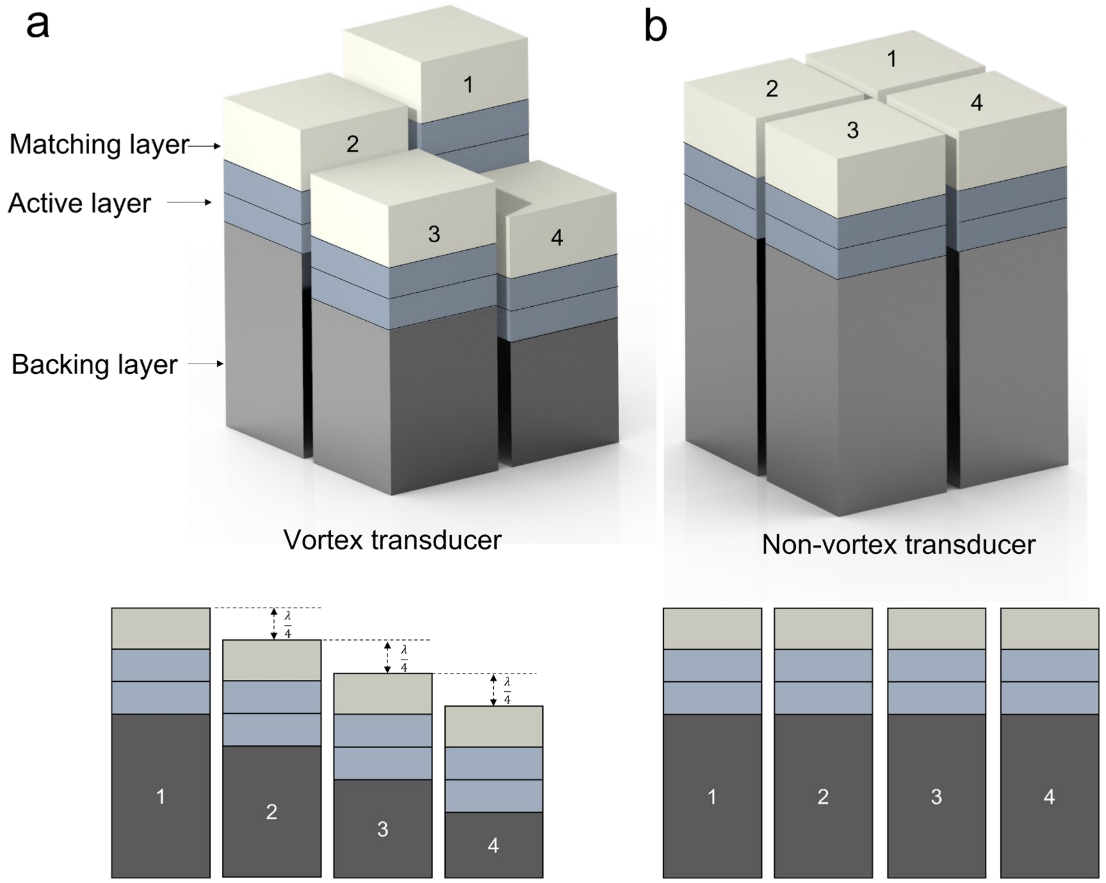


**Supplementary Fig. 1 | The design of vortex and nonvortex transducer. a.** The vortex transducer has four individual stack transducers, and each stack is composed of a matching layer on the top, an active layer in the middle, and a backing layer on the bottom. The backing layer height difference is controlled to be λ/4 to achieve the acoustic phase delay between adjoining transducers as π/2. **b.** The nonvortex transducer has a similar structure as the vortex transducer, but the backing layers have the same height so that there is no acoustic phase delay between adjacent transducers.

The multilayer design makes an electrically parallel and mechanically serial connection of stacked piezoelectric plates, resulting in a high ultrasonic transmitter efficiency while having a lower electrical impedance and a relatively low driving voltage with more significant strain. In our earlier forward-viewing sonothrombolysis transducer investigations, this transducer design was shown to be transmission efficient and generate substantial acoustic energy[1,2].

Moreover, unlike the mechanical phase delay caused by the physical shifting of the transducer surfaces, the electronic phase delay may be used to correctly adjust the emitting acoustic phase of each transducer as an alternate strategy for the formation of vortex ultrasound. For example, an alternative design of the ultrasound transducer array is a transducer array with a flat forward-viewing surface similar to that of a nonvortex transducer (Supplementary Fig. 1b) and proper electrical phase delay (π/2 between the neighboring transducers) in the A.C. voltage input on each transducer. Either analog all-pass filters or digital control circuits may be used to implement the electrical phase delay successfully[3,4]. Adjusting the electrical phase delay during the calibration process of the transducer array allows for achieving a symmetric pressure field. The rotation of the helical wavefront of vortex ultrasound can be controlled by altering the phase difference between the neighboring transducers, which allows for the switching of shear stress direction in the blood clot and may further enhance the lysis efficiency. In addition, the electrical control technology eliminates the need for perfect alignment of the front viewing surface of the transducers, which would otherwise be required.

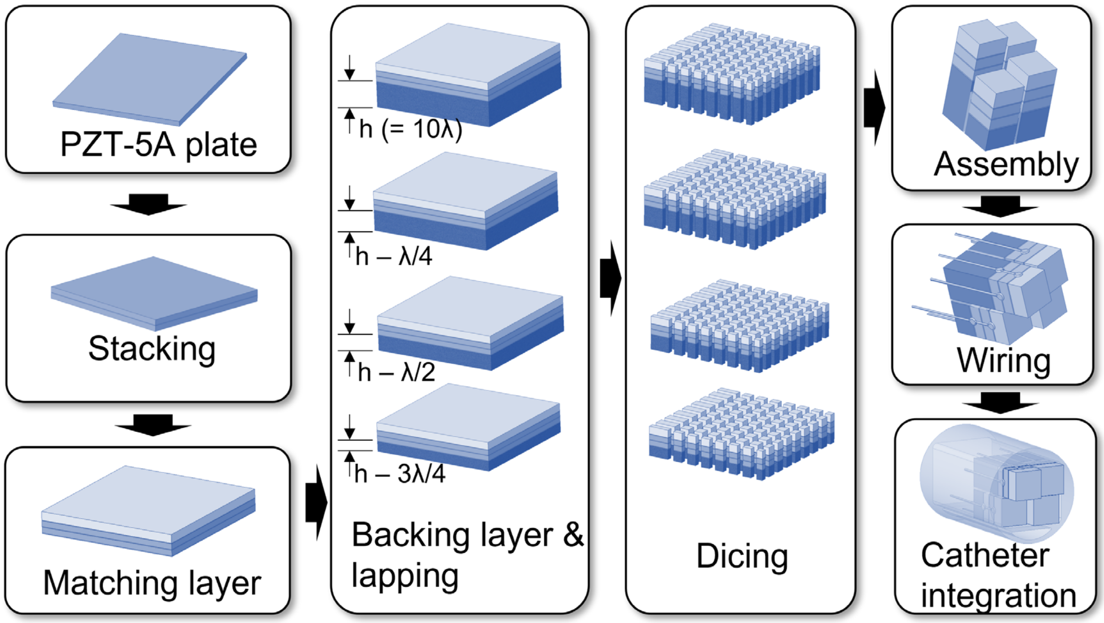


**Supplementary Fig. 2 |** The fabrication process of vortex transducer. The PZT-5A plate was first stacked and bonded together with an E-solder adhesive. The matching layer was lapped down to the desired thickness (h=λ/4) and then bonded onto the top of piezo stacks. The backing layer was lapped down to the different thicknesses with λ/4 height difference and then bonded onto the bottom of piezo stacks. The bonded stack was diced into individual transducers and assembled with epoxy. The coaxial cable was attached to the transducer electrodes for wire connection. Finally, the fabricated vortex transducer was integrated into a 9 Fr catheter.

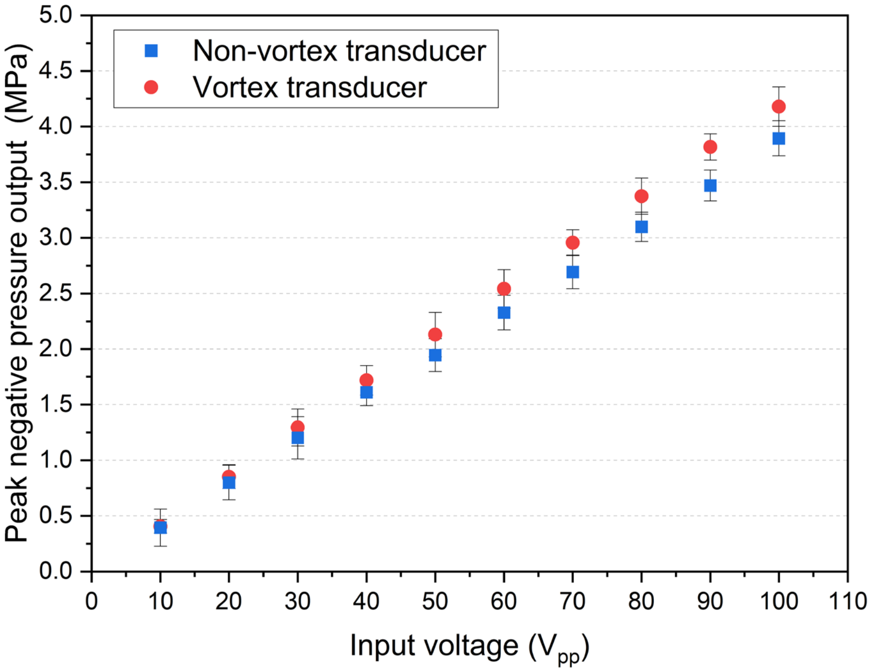


**Supplementary Fig. 3 |** The measured peak negative pressure of vortex and nonvortex transducers. With an input voltage of 100 $V_{pp}$, the transducer can reach a peak negative pressure of about 4 MPa.

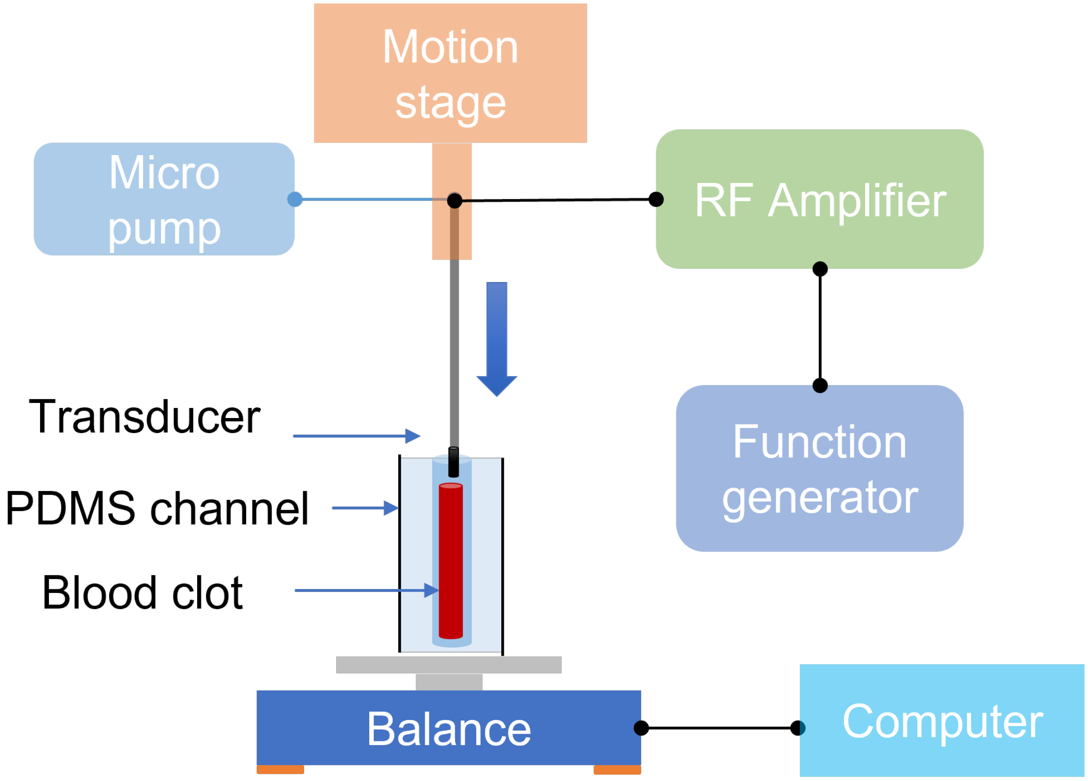


**Supplementary Fig. 4 |** The in vitro experimental setup. The acute blood clot was formed inside the PDMS channel, and the balance was used to measure the push-through force of the sonothrombolysis process. The transducer was powered by the RF amplifier with an input signal generated by a function generator. The position of the transducer was controlled by the motion stage with different feed-in speeds during the treatment. The micropump was used to inject the microbubbles solution for cavitation.

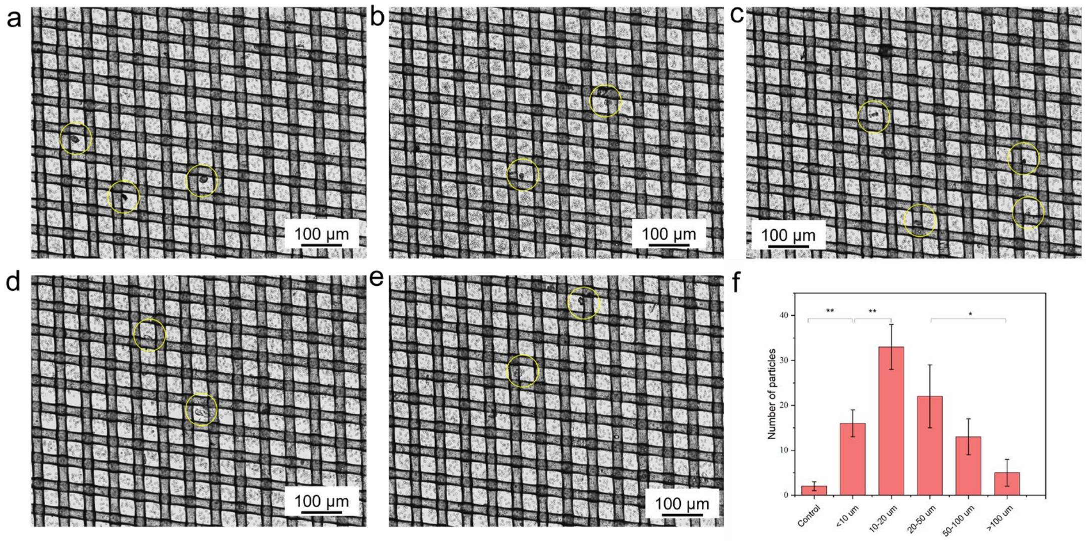


**Supplementary Fig. 5 |** The results of clot debris study. **a, b, c, d, e.** Clot debris collected after the treatment was examined under the microscope. The size of the mesh is about 50 µm. **f.** The size distribution of the clot debris (* $p<0.05$, ** $p<0.01$, $n=3$). Most clot debris particles measured less than 100 µm in diameter, indicating a minimal probability of dangerous embolus formation.

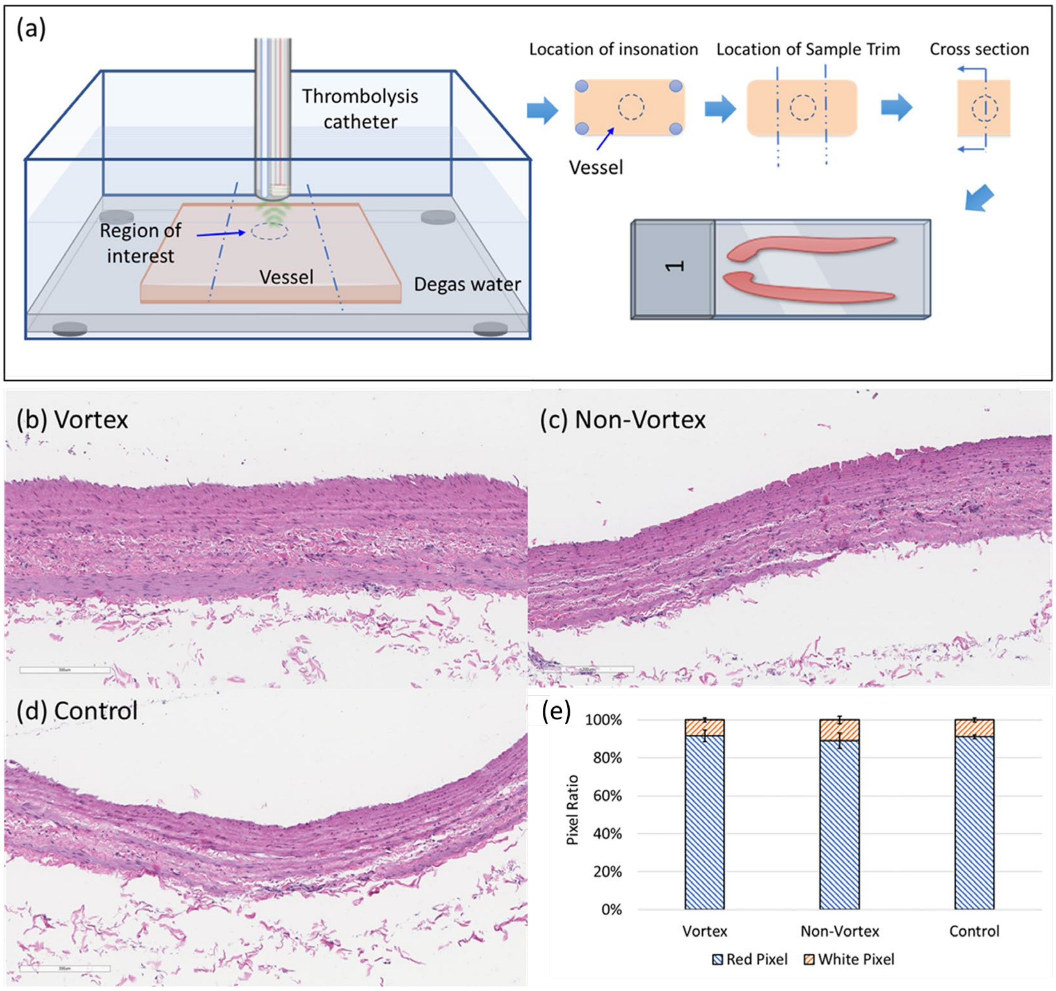


**Supplementary Fig. 6 |** The histology results of the bovine blood vessel wall cross-sections after operations of the vortex, nonvortex, and control groups. **a.** The schematic view of the experimental setup. **b.** The histology results of the vortex ultrasound treatment group. **c.** The histology results of the nonvortex ultrasound treatment group. **d.** The histology results of the control group. There was no macroscope damage seen on any vessel samples. Specifically, based on the histology findings, the orientation or spacing of endothelial cells along the arteries did not alter. **e.** The pixel ratios of vortex, nonvortex, and control group. According to the ratios of red and white pixels, there was no significant change between the three groups, indicating that neither nonvortex nor vortex transducers caused any harm to the vessel walls.

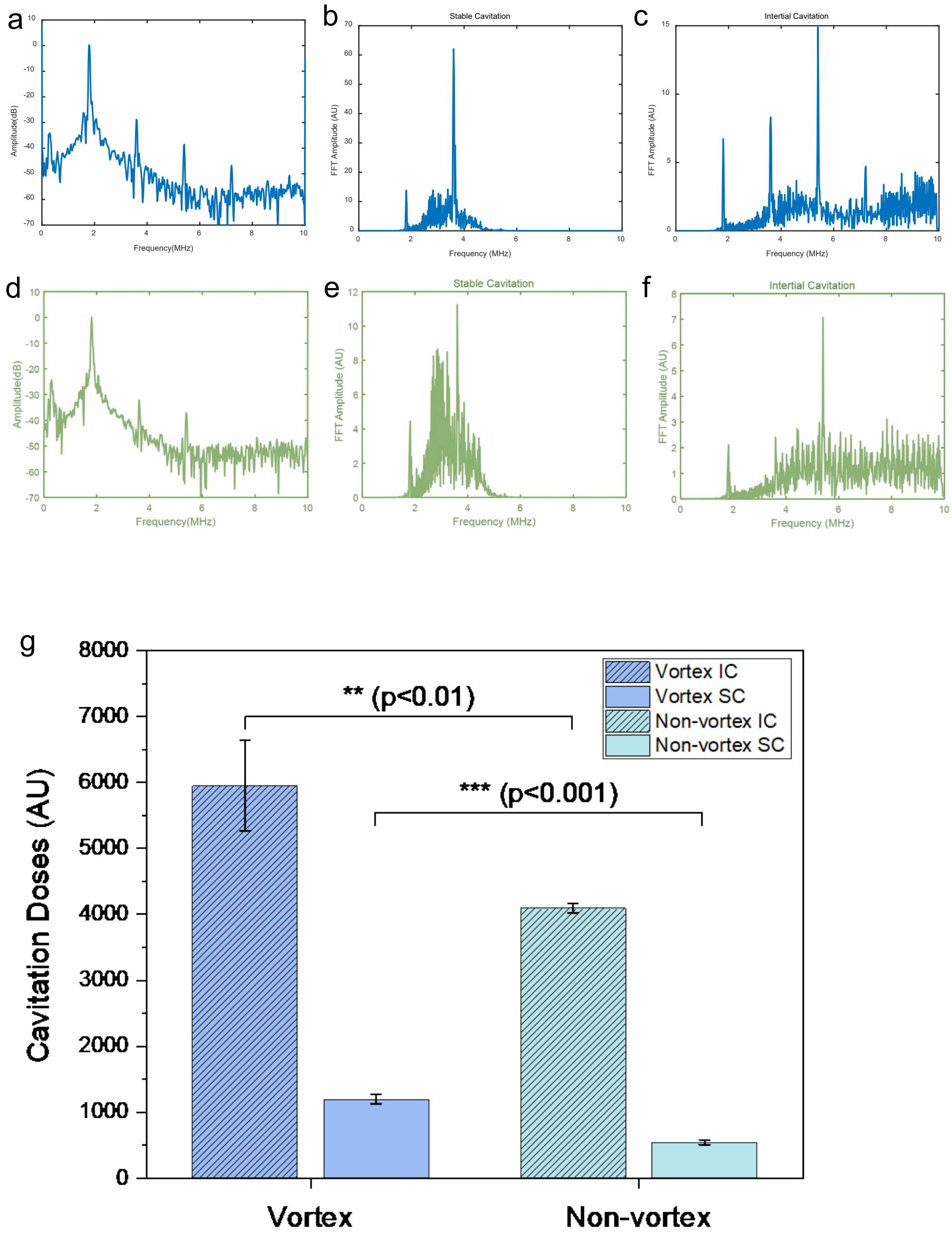


**Supplementary Fig. 7 |** The microbubbles cavitation comparison for vortex ultrasound and non-vortex ultrasound. **a**. Frequency spectrum of vortex ultrasound. **b**. Stable cavitation signal of vortex ultrasound. **c.** Inertia cavitation signal of vortex ultrasound. **d**. Frequency spectrum of non-vortex ultrasound. **e**. Stable cavitation signal of non-vortex ultrasound. **f.** Inertia cavitation signal of non-vortex ultrasound. **g**. The stable and inertia cavitation doses comparison between vortex and non-vortex ultrasound. The stable cavitation dose was calculated based on the area under the curve of the second harmonic ($2f_0 \pm 0.5f_0$) of the frequency spectrum (**a, d**) of the RF data. The inertial cavitation dose was calculated based on the area under the curve of the broadband noise of the 3rd–5th harmonics, wherein the fundamental harmonics ($3f_0 \pm 0.2f_0$, $4f_0 \pm 0.2f_0$, $5f_0 \pm 0.2f_0$) were subtracted from the whole AUC in this range. The ultrasound parameters: frequency: 1.8 MHz, duty cycle: 2.5%, 1 ms burst period, 45 cycles numbers.

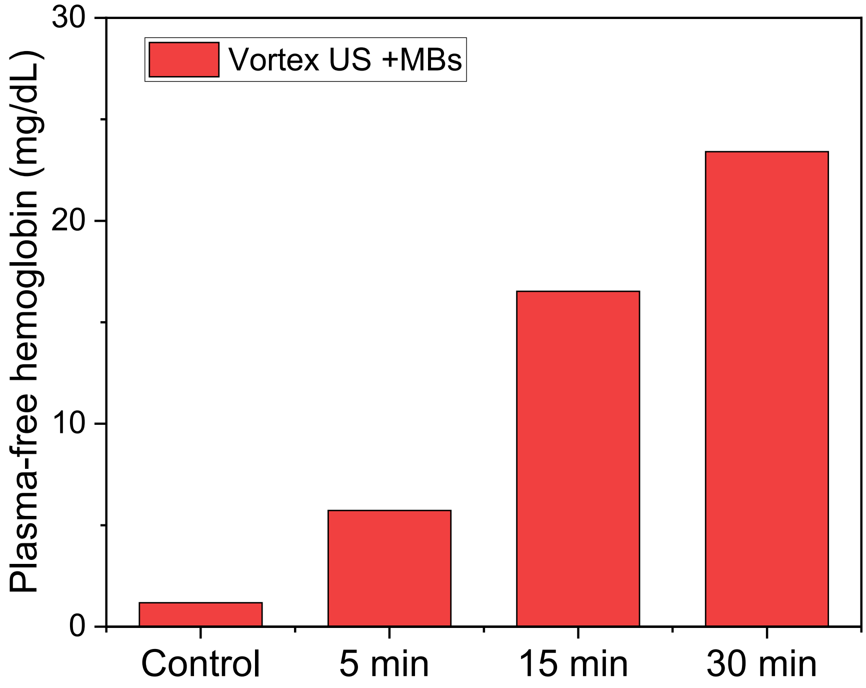


**Supplementary Fig. 8 |** The *in vitro* hemolysis test results under different treatment time (5 min, 15 min, 30 min) with vortex ultrasound (Frequency: 1.8 MHz, duty cycle: 7.5%, PRF: 10 kHz, input voltage: 80 $V_{pp}$) and microbubbles ($10^9$ MBs/mL) treatment. The control group was not treated with ultrasound or microbubbles.

**Movie S1 (separate file)**. The effectiveness of the vortex transducer in treating cerebral venous sinus thrombosis was evaluated with the use of an in vitro cerebral venous sinus 3D phantom flow model having an average sinus diameter of 10 millimeters. After only eight minutes of therapy with a vortex ultrasound transducer, the previously completely blocked blood artery was shown to have been successfully recanalized.
